# Topoisomerase II poisons inhibit vertebrate DNA replication through distinct mechanisms

**DOI:** 10.1101/2021.10.12.464107

**Authors:** Sabrina X. Van Ravenstein, Kavi P. Mehta, Tamar Kavlashvili, Jo Ann Byl, Runxiang Zhao, Neil Osheroff, David Cortez, James M. Dewar

## Abstract

Topoisomerase II (Top2) unlinks chromosomes during vertebrate DNA replication. Top2 ‘poisons’ are widely-used chemotherapeutics that stabilize Top2 complexes on DNA, leading to cytotoxic DNA breaks. However, it is unclear how these drugs affect DNA replication, which is a major target of Top2 poisons. Using *Xenopus* egg extracts, we show that the Top2 poisons etoposide and doxorubicin both inhibit DNA replication through different mechanisms. Etoposide induces Top2-dependent DNA breaks and induces Top2-dependent fork stalling by trapping Top2 behind replication forks. In contrast, doxorubicin does not lead to appreciable break formation and instead intercalates into parental DNA to inhibit replication fork progression. In human cells, etoposide stalls replication forks in a Top2-dependent manner, while doxorubicin stalls forks independently of Top2. However, both drugs exhibit Top2-dependent cytotoxicity. Thus, despite shared genetic requirements for cytotoxicity etoposide and doxorubicin inhibit DNA replication through distinct mechanisms.

## Introduction

Topoisomerases play essential roles during chromosome metabolism in all organisms (Keszthelyi et al. 2016; Pommier et al. 2016; Vann et al. 2021). Topoisomerases ordinarily unlink DNA by creating transient DNA breaks that are rapidly re-ligated. Topoisomerase ‘poisons’ are widely-used chemotherapeutics that prevent re-ligation of topoisomerase-induced breaks and trap topoisomerases on DNA, leading to cytotoxicity (Nitiss 2009; Pommier et al. 2016; Vann et al. 2021). Although the toxicity of topoisomerase poisons is largely due to their target topoisomerase, some topoisomerase poisons exert additional effects that may also contribute to their cytotoxicity (Doroshow et al. 2001; Koster et al. 2007; Coldwell et al. 2008; Pang et al. 2013; Yang et al. 2013). Additionally, the breaks induced by topoisomerase poisons appear to target specific aspects of chromosome metabolism (Markovits et al. 1987; Holm et al. 1989; Hsiang et al. 1989; Canela et al. 2017; Canela et al. 2019).

The most widely used topoisomerase poisons target topoisomerase II (Top2). This enzyme resolves supercoils as well as DNA intertwines between sister chromatids (‘catenanes’ and ‘pre-catenanes’) and DNA loops by creating double-strand DNA breaks through which another DNA strand is passed (Keszthelyi et al. 2016; Pommier et al. 2016; Vann et al. 2021). Break formation by type II topoisomerases involves formation of a covalent bond between catalytic tyrosine residues on the enzyme and 5’ DNA ends to create a specialized DNA-protein crosslink called a ‘cleavage complex’. Although double-strand DNA breaks are thought to be the major cause of cytotoxicity by Top2 poisons, most Top2 complexes trapped by Top2 poisons do not generate double-strand breaks (Muslimović et al. 2009; Canela et al. 2019). Thus, a major effect of Top2 poisons is interference with Top2 activity, rather than formation of double-strand DNA breaks.

Vertebrates contain two isoforms of Top2, denoted α and β (Austin et al. 2018). Top2α is essential for DNA replication and mitosis, where its essential role is to unlink newly replicated chromosomes (Grue et al. 1998; Linka et al. 2007). Top2β cannot carry out this function and instead is involved in transcription and DNA loop formation (Linka et al. 2007; Uusküla-Reimand et al. 2016; Canela et al. 2017). Importantly Top2α is essential for cell viability while Top2β is not, indicating that the essential functions of Top2β can also be fulfilled buy Top2α (Grue et al. 1998; Linka et al. 2007; Uusküla-Reimand et al. 2016; Canela et al. 2017). Both isoforms of Top2 are targeted by chemotherapeutics such as etoposide and doxorubicin, which are widely used treatments for lung, breast, and ovarian cancers (Murai 2017; Waqar and Morgensztern 2017). Importantly, much of the cytotoxic effect of Top2 poisons is due to targeting Topα during DNA replication (Markovits et al. 1987; Burgess et al. 2008; Fan et al. 2008; Tammaro et al. 2013; Pommier et al. 2016). During vertebrate DNA replication Top2α resolves pre-catenanes throughout replication to carry out the majority of sister chromatid unlinking (Heintzman et al. 2019). Resolution of pre-catenanes by Top2α is also important to prevent accumulation of topological stress that would otherwise stall replication forks during the final stages of DNA synthesis (Heintzman et al. 2019). After completion of DNA synthesis, any catenanes not protected by cohesin are also removed by Top2α (Sumara et al. 2002; Farcas et al. 2011). Since Top2α works throughout DNA replication, all stages of DNA replication are potentially susceptible to Top2 poisons.

Top2 poisons are potent inhibitors of DNA synthesis (Zellweger et al. 2015; Pommier et al. 2016). One model to explain this inhibition is that Top2 poisons trap Top2 ahead replication forks on DNA supercoils, creating a cytotoxic lesion when the replisome collides with the trapped topoisomerase (Figure S1Aii), similar to how topoisomerase I poisons function (Strumberg et al. 2000; Yan et al. 2016). However, this model has not been directly tested and Top2α primarily acts behind forks during vertebrate replication (Lucas et al. 2001; Heintzman et al. 2019), suggesting that replication inhibition must occur through another mechanism.

Etoposide was reported to indirectly inhibit DNA replication by inducing single-strand breaks that lead to activation of the ATR kinase and subsequent inhibition of replication initiation (Figure S1Aiii) (Costanzo et al. 2003). However, this cannot explain inhibition of fork progression (Zellweger et al. 2015) after initiation has taken place. Doxorubicin was also reported to indirectly inhibit replication initiation by preventing nuclear envelope formation (Krasinska and Fisher 2009). However, this activity was independent of Top2 and likely reflected doxorubicin’s intercalative properties (Yang et al. 2015). Thus, although Top2 poisons inhibit replication fork progression it is unclear how this occurs.

In this study, we determined the effects of the Top2 poisons etoposide and doxorubicin during vertebrate DNA replication using *Xenopus* egg extracts. We found that both etoposide and doxorubicin directly inhibit DNA synthesis, but etoposide behaves as a canonical Top2 poison during DNA replication while surprisingly doxorubicin does not. We show that etoposide traps Top2α behind forks to inhibit replication fork progression, suggesting that it stalls replication forks by interfering with resolution of topological stress. Strikingly, doxorubicin inhibits replication independently of Top2α by acting as a DNA intercalator. We corroborated our results by analyzing DNA replication in mammalian cells, which showed that etoposide inhibits replication in a Top2α-dependent manner, while doxorubicin inhibits replication independent of Top2α, as observed using *Xenopus* egg extracts. Thus, our data show that etoposide and doxorubicin directly inhibit replication fork progression through different mechanisms despite shared genetic requirements for toxicity.

## Results

### Differential effects of topoisomerase II poisons on DNA replication

We first tested how etoposide and doxorubicin impact DNA replication in the completely-soluble *Xenopus* egg extract system (Walter et al. 1998). In this system, replication typically initiates from a single origin per plasmid and DNA synthesis is monitored by the incorporation of radioactive nucleotides (Fig. 1A). We performed a titration series of each drug and found that a wide range of concentrations inhibited DNA replication in a dose-dependent manner (Figs 1B-D and Supplemental Fig. S1B-G). Surprisingly, inhibition of replication by etoposide was not dependent on ATR signaling, and thus was distinct from the previously described mechanism in *Xenopus* egg extracts (Costanzo et al. 2003) (Supplemental Fig. S2A-E). Furthermore, replication in this extract system (Walter et al. 1998) does not involve a nuclear envelope, so inhibition of replication by doxorubicin could not be due to impaired nuclear envelope assembly, as previously described in *Xenopus* egg extracts (Krasinska and Fisher 2009). Thus, inhibition of DNA replication by etoposide and doxorubicin can occur by different mechanisms to those previously described.

**Figure 1.**
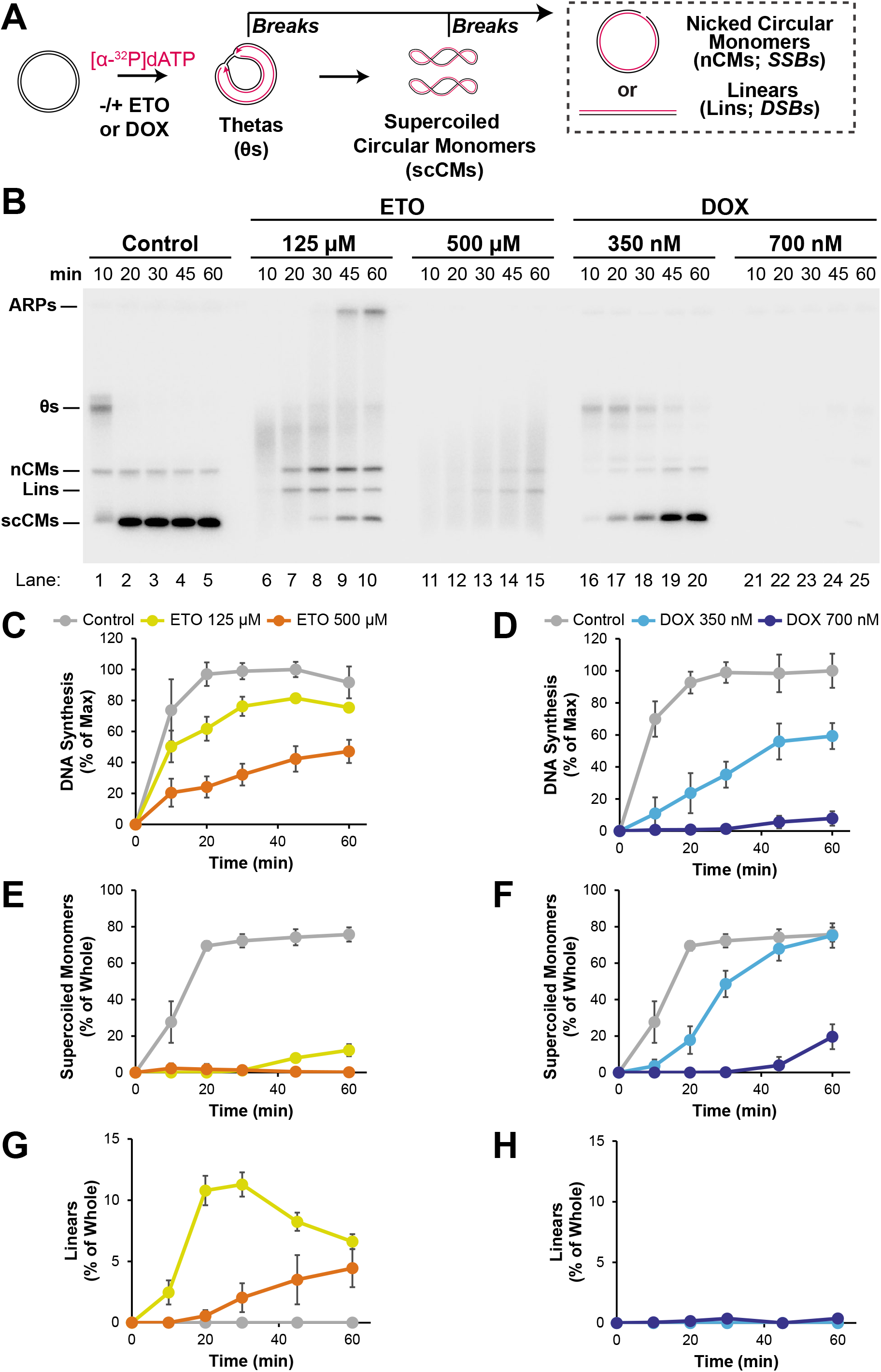
Top2 poisons exert different effects on DNA replication. *(A)* Plasmid DNA was replicated using *Xenopus* egg extracts in the presence of [α-^32^P]dATP to label newly-synthesized DNA strands. Extracts were treated with either DMSO (Control), 125 μM etoposide, 500 μM etoposide, 350 nM doxorubicin, or 700 nM doxorubicin at the onset of DNA replication. *(B)* Samples from (A) were separated on a native agarose gel and visualized by autoradiography (See also Figure S1B,E). *(C)* Quantification of total DNA synthesis from (B) lanes 1-15. Mean ± SD, n=3 independent experiments. *(D)* Quantification of total DNA synthesis from (B) lanes 1-5, 16-25. Mean ± SD, n=3 independent experiments. *(E)* Quantification of supercoiled circular monomers (‘scCMs’) from (B) lanes 1-15. Mean ± SD, n=3 independent experiments. *(F)* Quantification of scCMs from (B) lanes 1-5, 16-25. Mean ± SD, n=3 independent experiments. *(G)* Quantification of linears (‘lins’) from (B) lanes 1-15. Mean ± SD, n=3 independent experiments. *(H)* Quantification of lins from (B) lanes 1-5, 16-25. Mean ± SD, n=3 independent experiments.

Intriguingly, the DNA structures formed by etoposide and doxorubicin treatment were very different. In the presence of etoposide we observed linear molecules and aberrant replication products (Fig. 1B, lanes 6-15, Lins and ARPs), consistent with formation of DNA double-strand breaks (‘DSBs’). In contrast doxorubicin treatment did not alter the DNA structures formed (Fig. 1B, lanes 16-25). To quantify this difference we compared 125 μM and 500 μM etoposide and 350 nM and 700 nM doxorubicin, which led to sub-maximal inhibition of DNA replication by each drug (Fig. 1C-D and Supplemental Fig. S1B-G). Using these concentrations, we measured supercoiled circular monomers (Fig. 1A, scCMs), which are the final replication product in this system. We found that even though 125 μM etoposide was low enough that total DNA synthesis was largely unaffected (Fig. 1C) the formation of scCMs was essentially blocked (Fig. 1B, lanes 6-10, 1E, ETO 125 μM). In contrast, although 350 nM doxorubicin more potently inhibited DNA synthesis (Fig. 1D) it still resulted in the formation of scCMs by most molecules (Fig. 1B, lanes 16-20, 1F). Higher concentrations of etoposide and doxorubicin also showed that even though 700 nM doxorubicin exhibited stronger inhibition of DNA synthesis than 500 μM etoposide (Fig. 1C-D) it did not inhibit scCM formation as strongly as etoposide (Fig. 1B, lanes 11-15 and 21-25, E-F, and Supplemental Fig. S2F). Thus, at comparable levels of replication inhibition etoposide inhibits scCM formation more potently than doxorubicin. Linear molecules were also much more abundant in the presence of etoposide than doxorubicin (Fig. 1G-H), which suggested that the more potent inhibition of scCM formation by etoposide was due to an increased ability to induce DNA breaks. Overall, we conclude that etoposide and doxorubicin have different effects on vertebrate DNA replication and may result in different frequencies of DNA break formation.

### Etoposide, but not doxorubicin, acts as a canonical Top2α poison during DNA replication

Topoisomerase II (Top2) poisons stabilize Top2 cleavage complexes, which results in accumulation of DNA-protein cross-links (‘DPCs’) as well as DNA DSBs (Pommier et al. 2016; Vann et al. 2021). Given the apparent lack of DSBs following doxorubicin treatment (above) we wanted to directly examine the ability of etoposide and doxorubicin to induce DSBs and DPCs during DNA replication. To this end, we performed replication reactions in the presence of etoposide and doxorubicin, purified replication intermediates, and separated restriction digested products on a native agarose gel (Fig. 2A). Under these conditions, fully replicated intact molecules should produce linear molecules the size of the full-length plasmid (Fig. 2A, Lins, 3148 bp) and DSBs should form smaller fragments (Fig. 2A, ‘Breaks’, <3148 bp). In the vehicle control we did not readily detect breaks (Fig. 2B, lanes 1-5, 2C). Following etoposide treatment, breaks were readily detected, as expected (Fig. 2B, lanes 6-10, 2C). Strikingly, breaks were not readily detected in the doxorubicin condition (Fig. 2B, lanes 11-15) and the level was indistinguishable from background (Fig. 2C). The concentrations of etoposide and doxorubicin used in these experiments inhibited formation of fully replicated linear molecules to a similar extent (Fig. 2D) indicating comparable levels of replication inhibition. Thus, inhibition of replication by etoposide readily induces DSBs, while doxorubicin does not.

**Figure 2.**
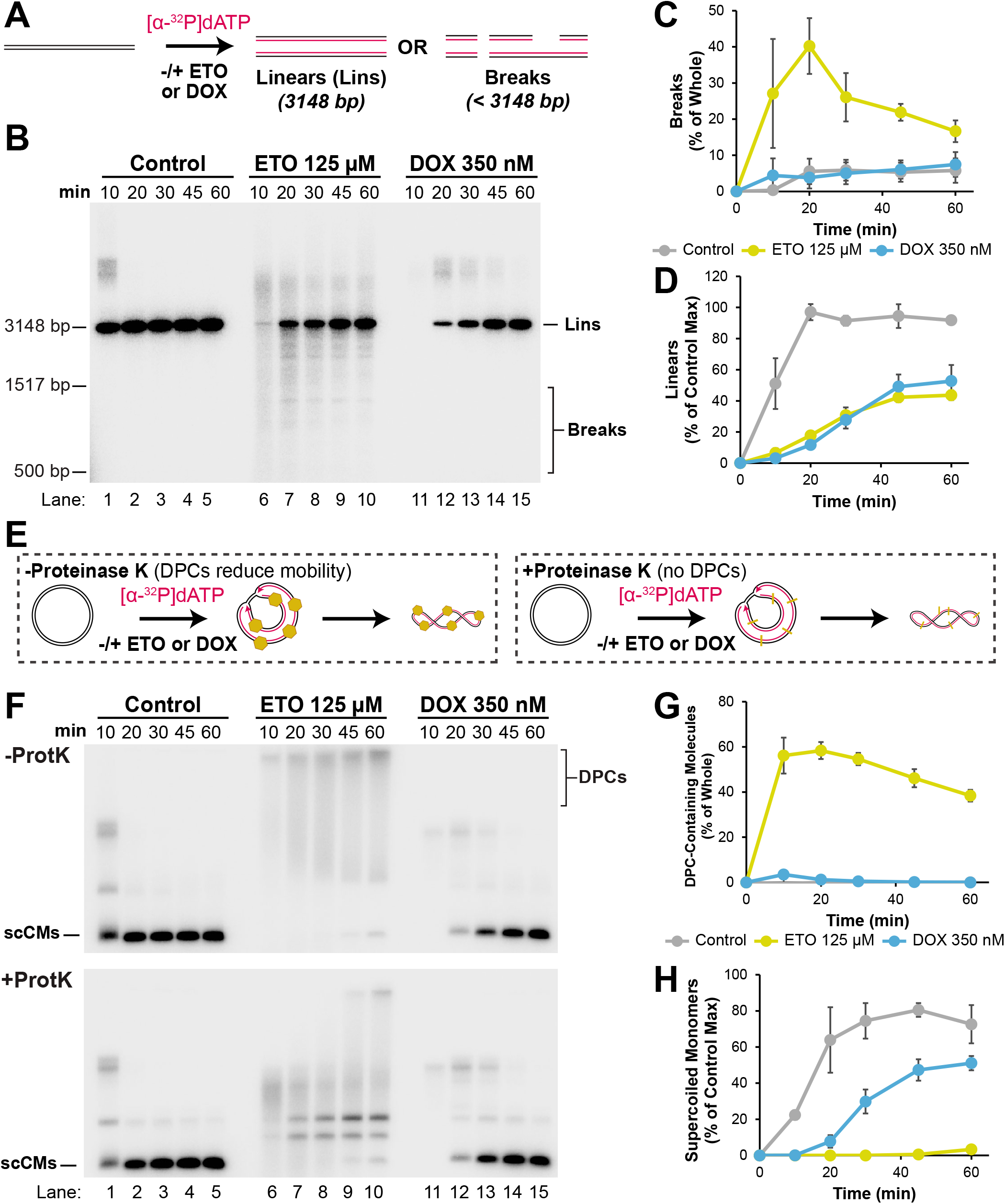
Etoposide, but not doxorubicin, creates breaks and DNA-protein crosslinks during DNA replication. *(A)* Plasmid DNA was replicated in the presence of [α-^32^P]dATP and vehicle control, etoposide, or doxorubicin. Purified DNA intermediates were digested with XmnI, which cuts the plasmid template once, to allow for identification of DNA breaks. *(B)* Samples from (A) were separated on a native agarose gel and visualized by autoradiography. *(C)* Quantification of DNA breaks from (B). Mean ± SD, n=3 independent experiments. *(D)* Quantification of linears (Lins)from (B). Mean ± SD, n=3 independent experiments. *(E)* Plasmid DNA was replicated as in (A). Samples were split and either treated with (‘+Prot K’) or without (‘-Prot K’) proteinase K. *(F)* Samples from (E) were separated on native agarose gels and visualized by autoradiography. *(G)* Quantification of DPC-containing molecules from (F). Mean ± SD, n=3 independent experiments. *(H)* Quantification of the amount of supercoiled circular monomers (‘scCMs’) from ‘+ProtK’ samples from (F). Mean ± SD, n=3 independent experiments.

Next, we wanted to determine the ability of etoposide and doxorubicin to induce DPCs during DNA replication. We replicated DNA in the presence of etoposide or doxorubicin and either added or omitted proteinase K to detect DPCs, as previously described (Duxin et al. 2014) (Fig. 2E). In the vehicle control proteinase K had essentially no effect on migration of replication intermediates (Fig. 2F, lanes 1-5, -ProtK and +ProtK) indicating that DPCs were not formed. Following etoposide treatment, DPC-containing molecules ran as high molecular weight smears on native agarose gels in the absence of proteinase K (Fig. 2F, -ProtK) and these were largely resolved by proteinase K treatment (Fig. 2F, +ProtK) indicating that DPCs were formed. In contrast, the DNA structures formed by doxorubicin treatment were essentially unaffected by proteinase K (Fig. 2F, lanes 11-15) indicating few or no DPCs. Quantification of DPCs showed that etoposide dramatically induced DPC formation while the level of DPC induction by doxorubicin was indistinguishable from background (Fig. 2G). Quantification of scCMs revealed that doxorubicin still impacted DNA replication, even though DPCs were not readily detected (Fig. 2H). Thus, during DNA replication etoposide readily induces DPCs while doxorubicin does not, as observed for DSBs (above).

Our experiments show that etoposide and doxorubicin exhibit different effects on DNA replication. Since both drugs induce cell killing in a Top2-dependent manner we wanted to know whether the different effects they exerted on replication depended on Top2. To address this, we immunodepleted the main isoform of Top2, Top2α (Gaggioli et al. 2013; Wühr et al. 2014), from extracts and monitored replication inhibition after etoposide and doxorubicin treatment (Fig. 3A). Following mock immunodepletion the final replication products were scCMs (Fig. 3B, lanes 1-5,) and formation of these species was inhibited by etoposide and doxorubicin (Fig. 3B, lanes 6-15, 3C) as previously observed (Fig. 1). Following depletion of Top2α, the final replication products were catenated dimers since there was no Top2α to carry out unlinking (Fig. 3B, lanes 16-30, ‘Cats’). Accumulation of these species was unaffected by etoposide (Fig. 3B, lanes 16-25, 3D), consistent with the effects of etoposide being dependent upon Top2. In contrast, doxorubicin still inhibited formation of catenanes in Top2α-depleted extracts (Fig. 3B, lanes 16-20 and 26-30, 3D). Moreover, the extent of inhibition was similar to the inhibition of supercoiled circular monomers following mock immunodepletion (Fig. 3C-D), suggesting the effect of doxorubicin were largely independent of Top2. Etoposide and doxorubicin also exhibited the same behavior when we monitored their effects on formation of breaks and DPCs during replication in the presence or absence of human Top2α (Supplemental Fig. S3). Thus, these data show that etoposide inhibits replication in a Top2α-dependent manner while doxorubicin does not.

**Figure 3.**
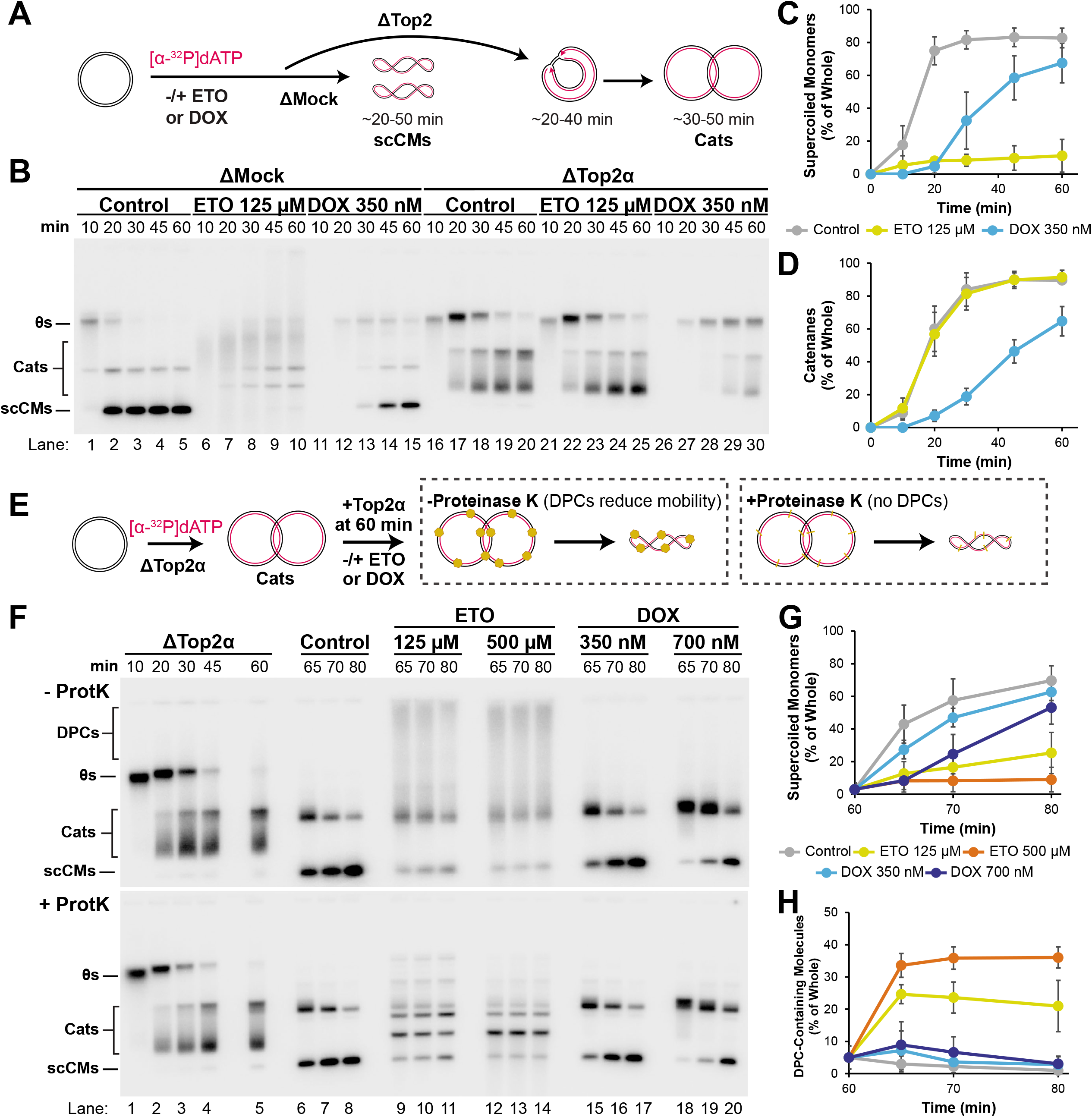
Etoposide and doxorubicin inhibit DNA replication through different mechanism. *(A)* Plasmid DNA was replicated in Mock- or Top2α-immunodepleted extracts in the presence or absence of vehicle control, etoposide, or doxorubicin. In mock-depleted extracts (ΔMock), the final replication product are supercoiled circular monomers (‘scCMs’). In Top2α-depleted extracts (ΔTop2α), the final replication product are catenanes (‘Cats’) because Top2α is required for DNA unlinking. *(B)* Samples from (A) were separated on a native agarose gel and visualized by autoradiography. *(C)* Quantification of scCMs from (B) lanes 1-15. Mean ± SD, n=3 independent experiments. *(D)* Quantification of Cats from (B) lanes 16-30. Mean ± SD, n=3 independent experiments. *(E)* Plasmid DNA was replicated in Mock- or Top2α-immunodepleted extracts. At 60 minutes, undepleted extract was added back to add to supply Top2α and trigger decatenation. Vehicle, etoposide, or doxorubicin were also added at 60 minutes. Samples were treated either with or without Proteinase K (as in Figure 2E). *(F)* Samples from (E) were separated on native agarose gels and visualized by autoradiography. *(G)* Quantification of scCMs from + ProtK gel in (F) lanes 6-20. Mean ± SD, n=3 independent experiments. *(H)* Quantification of DPC Containing Molecules from (F) lanes 6-20. Mean ± SD, n=3 independent experiments.

We reasoned that the inhibition of DNA replication by doxorubicin might prevent formation of catenanes, which are the substrate for Top2α during replication in *Xenopus* egg extracts (Heintzman et al. 2019) and thus prevent formation of breaks and DPCs by depriving Top2α of its substrate. To test this possibility, we developed a scheme to test the ability of etoposide and doxorubicin to induce DPCs under conditions where a fixed number of catenanes were present at the start (Fig 3E). Top2α was immunodepleted from extracts and DNA was replicated for 60 minutes to produce primarily catenated dimers (Fig. 3E and 3F, lanes 1-5). A small quantity of undepleted extract was then added back to provide Top2 activity in the absence or presence of etoposide or doxorubicin (Fig. 3E) and DPCs were then detected (as above). In the vehicle control, catenated dimers were rapidly resolved to scCMs and DPCs were not detected (Fig. 3F, lanes 6-8, 3G and 3H). In contrast, etoposide treatment strongly inhibited formation of supercoiled circular monomers and DPCs were readily detected (Fig. 3F, lanes 9-14, 3G and 3H), as observed previously (Fig. 2E-H). Importantly, doxorubicin still did not appreciably produce DPCs (Fig. 3F, lanes 15-20, 3G and 3H), as observed previously (Fig. 2E-H). Strikingly the extent of scCM formation formed following 350 nM doxorubicin treatment was almost indistinguishable from the vehicle control (Fig. 3G). Thus, the lack of DPC formation following doxorubicin treatment could not be attributed to either lack of catenanes or lack of Top2 activity. Taken together, these data show that etoposide behaves as a canonical Top2α poison during DNA replication while doxorubicin does not.

### Etoposide stalls forks by trapping Top2α on newly-replicated DNA

Top2 poisons are thought to inhibit replication by creating breaks ahead of replication forks, leading to fork collapse (Supplemental Fig. S1ii) or by creating breaks behind forks that activate the ATR checkpoint and inhibit DNA synthesis (Supplemental Fig. S1iii). Inhibition of replication by etoposide did not require the ATR checkpoint (Supplemental Fig. S2A-E) suggesting that the inhibition of replication was instead due to fork collapse. To test this, we replicated DNA in the absence or presence of etoposide, then recovered chromatin-bound proteins and analyzed their abundance by Western blotting (Fig. 4A). As expected, etoposide treatment led to accumulation of chromatin-bound Top2α (Fig. 4B, lanes 5-8 and 13-16, 4C). Top2α included higher molecular weight species, which we speculate arise from sumoylation or ubiquitylation of Top2 cleavage complexes (Sun et al. 2020). If etoposide caused fork collapse, we would expect to see replisome components dissociate prematurely due to translocation off the DNA template when a break was encountered (Supplemental Fig. S1Aii). However, in contrast to this prediction we observed that the core replicative helicase component CDC45 persisted on DNA (Fig. 4B, lanes 1-8, 4E). These data indicate that replication forks are stalled by etoposide rather than prematurely disassembled. We conclude that etoposide stalls replication fork progression.

**Figure 4.**
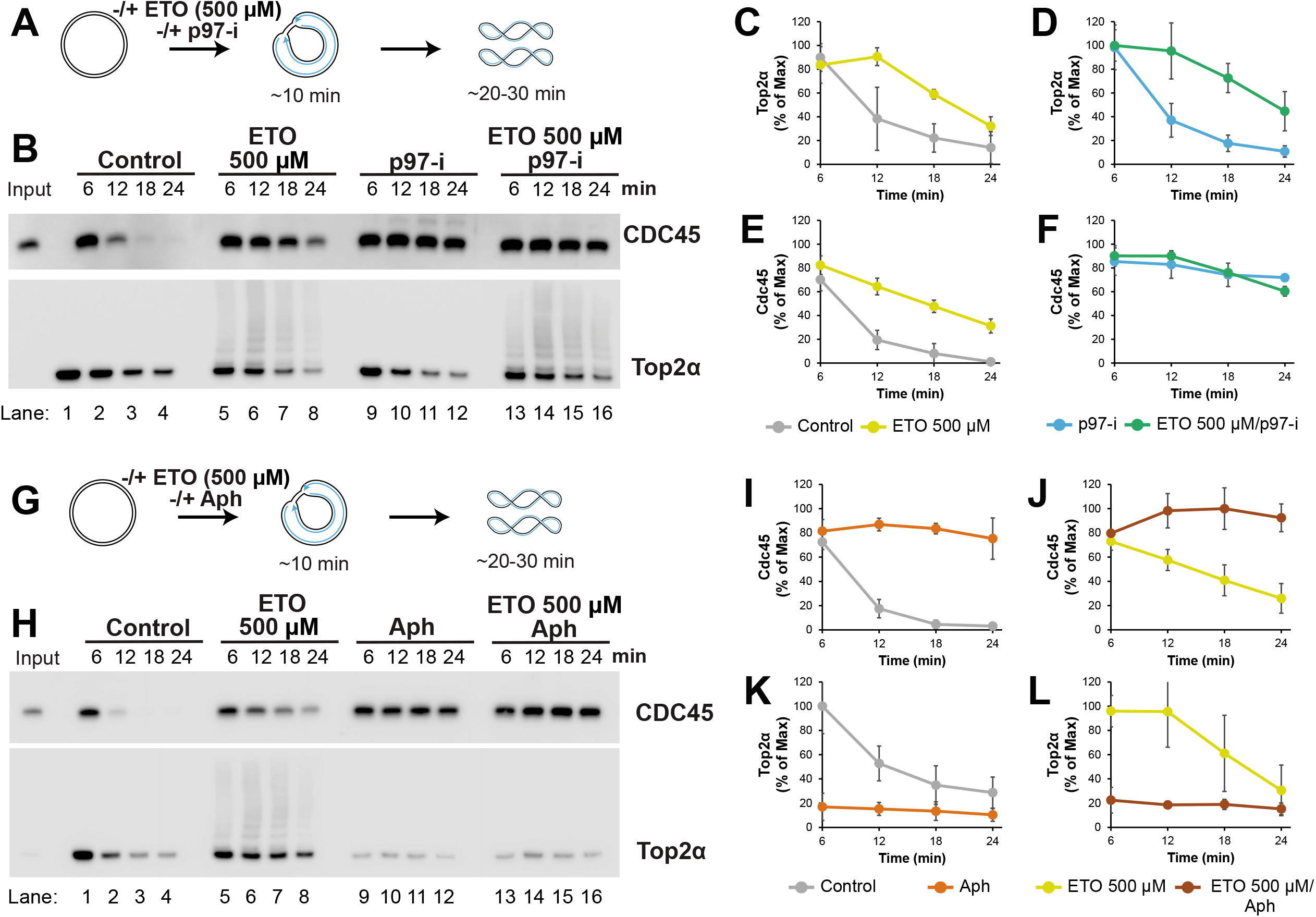
Etoposide stalls replication forks by trapping Top2α behind forks. *(A)* Plasmid DNA was replicated in the presence or absence of etoposide and p97-i. At different timepoints chromatin-bound proteins were recovered via plasmid pulldowns. *(B)* Proteins from (A) were recovered and analyzed via Western blotting. Input corresponds to 1% of total reaction. *(C)* Quantification of Top2α from (B) lanes 1-8. *(D)* Quantification of Top2α from (B) lanes 9-16. *(E)* Quantification of CDC45 from (B) lanes 1-8. *(F)* Quantification of CDC45 from (B) lanes 9-16. Mean ± SD, n=3 independent experiments for (C)-(F). *(G)* Plasmid DNA was replicated in the presence or absence of etoposide and aphidicolin (Aph). At different timepoints chromatin-bound proteins were recovered via plasmid pulldowns. *(H)* Proteins from (G) were recovered and analyzed via Western blotting. Input corresponds to 1% of total reaction. *(I)* Quantification of CDC45 from (H) lanes 1-4 and 9-12. *(J)* Quantification of CDC45 from (H) lanes 5-8 and 13-16. *(K)* Quantification of Top2α from (H) lanes 1-4 and 9-12. *(L)* Quantification of Top2α from (H) lanes 5-8 and 13-16. Mean ± SD, n=3 independent experiments for (I)-(L). See also Supplemental Fig. S4B-D.

CDC45 ultimately did dissociate from DNA in the presence of etoposide (Fig. 4B). We wanted to know whether this reflected passive unloading following encounter with a DNA break (Supplemental Fig. S1Aii) or the replisome unloading pathways that operate following completion of DNA synthesis (Dewar et al. 2017; Sonneville et al. 2017; Deng et al. 2019; Priego Moreno et al. 2019; Wu et al. 2019). To test this, we once again monitored protein binding in the absence or presence of etoposide but added included a small molecule inhibitor of p97 (‘p97-i’), which inhibits all known replisome unloading pathways in *Xenopus* egg extracts (Dewar et al. 2017; Deng et al. 2019) but does not block replisome dissociation when a break in the translocating strand is encountered (Vrtis et al. 2021). Etoposide still led to enrichment of Top2α on DNA in the presence of p97-i (Fig. 4B, lanes 1-8 and 9-16, 4D). However, p97-i strongly inhibited dissociation of CDC45 and this was essentially unaffected by etoposide (Fig. 4B, lanes 9-16, 4F). Thus, the replisome unloading observed in the presence of etoposide is due to replisome unloading at the end of DNA replication rather than passive unloading caused by encounter with a break (Vrtis et al. 2021).

We next wanted to address whether etoposide traps Top2α ahead of or behind replication forks. Although Top2α acts behind forks during vertebrate replication (Heintzman et al. 2019), it was reported that etoposide can trap Top2α ahead of forks, albeit infrequently (Lucas et al. 2001). To distinguish these possibilities, we monitored binding of Top2α to DNA in the presence or absence of the DNA polymerase inhibitor aphidicolin (Fig. 4G), which blocks DNA synthesis but not unwinding (Walter and Newport 2000), and thus prevents formation of catenanes but not supercoils (Supplemental Fig. S4A). Aphidicolin treatment caused CDC45 and MCM6 to persist on chromatin, consistent with fork stalling (Fig. 4H lanes 9-12, 4I, and Supplemental Fig. S4B-D) and this was also true in the presence of etoposide (Fig. 4H lanes 13-16, 4J, and Supplemental Fig. S4C,D). Aphidicolin also dramatically inhibited Top2α binding (Fig. 4H lanes 9-12, 4K) as previously described, consistent with Top2α acting behind forks (Heintzman et al. 2019). If etoposide trapped Top2α ahead of the forks, then etoposide should lead to a substantial increase in Top2α binding event in the presence of aphidicolin. Instead, we found that aphidicolin essentially blocked Top2α binding, even in the presence of etoposide (Fig. 4H, lanes 9-16, 4K and 4L). Thus, etoposide traps Top2α behind forks on catenanes.

We previously found that catalytically inactive Top2α, which would also be expected to be trapped behind forks, interfered with replication fork progression (Heintzman et al. 2019). To test whether Top2α complexes trapped behind forks by etoposide could interfere with Top2 activity, we tested whether etoposide could reproduce the termination-specific fork stalling that we previously observed following loss of Top2 activity (Heintzman et al. 2019). To this end, we monitored fork merger, which represents unwinding of the final stretch of DNA duplex (Supplemental Fig. S4E), in the presence of limiting concentrations of Top2α so that we could observe formation of replication products without DNA double-strand breaks (Supplemental Fig. S4F). Etoposide treatment inhibited fork merger (Supplemental Fig. S4F lanes 6-10, S4G) but had no effect on total DNA synthesis (Supplemental Fig. S4H). Thus, etoposide inhibited fork merger. Overall, our data show that etoposide stalls replication fork progression by trapping Top2α behind forks and this can be explained by inhibition of Top2α activity.

### Doxorubicin stalls replication by intercalating unreplicated DNA

Doxorubicin has two well-defined activities – it induces Top2-dependent DNA breaks and functions as a DNA intercalator (Yang et al. 2014). Since doxorubicin did not create DNA breaks (Figs. 2-3) we wanted to test whether its DNA intercalator activity was sufficient to explain the replication inhibition we observed (Fig. 1). To address this point, we compared the effects of doxorubicin to aclarubicin, a similar anthracycline drug that intercalates DNA but does create Top2-dependent DNA breaks (Pang et al. 2013; Yang et al. 2014). We first identified concentrations of each drug that inhibited total DNA synthesis to the same extent (Supplemental Fig. S5A) and found that 2 µM and 4µM aclarubicin inhibited replication to the same extent as 350 nM and 700 nM doxorubicin, respectively. The requirement for higher concentrations of aclarubicin is consistent with aclarubicin being less potent than doxorubicin (Pang et al. 2015). We then monitored the effect of these drugs at on formation of scCMS, which are the final products of replication (Fig. 5A). The formation and resolution of scCMs formed by 2 µM aclarubicin was indistinguishable from 350 nM doxorubicin (Fig. 5B, lanes 6-10 and 16-20,). Likewise, formation and resolution of scCMs formed by 4 µM aclarubicin was indistinguishable from 700 nM doxorubicin (Fig. 5B, lanes 11-15 and 21-25). Quantification of these data revealed that the degree of inhibition of scCM formation was essentially identical for equivalent concentrations of doxorubicin and aclarubicin (Fig. 5C, 2 µM Aclarubicin and 350 nM Doxorubicin, 4 µM Aclarubicin and 700 nM Doxorubicin). Thus, DNA the intercalation activity of doxorubicin is sufficient to explain its inhibition of DNA replication.

**Figure 5.**
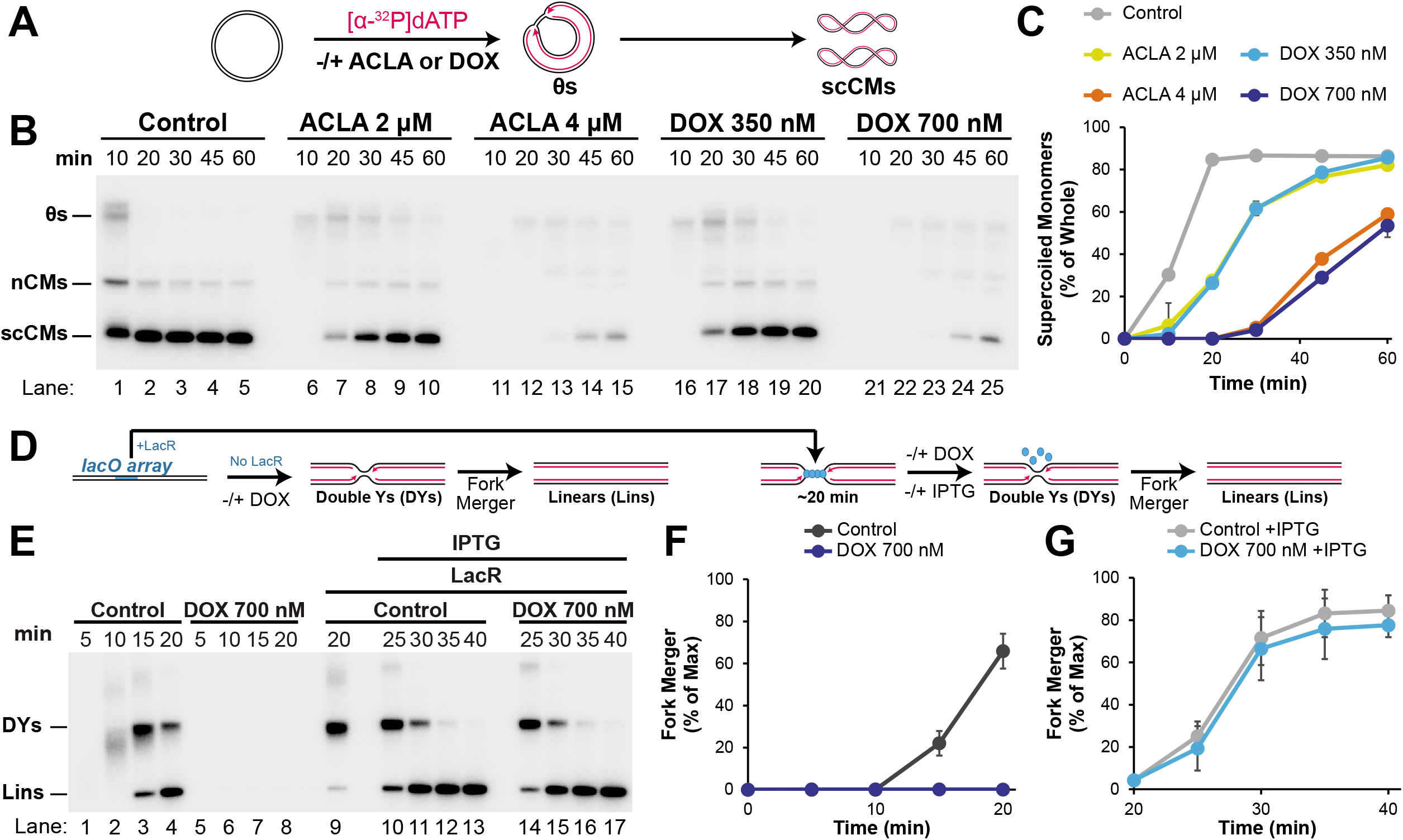
Doxorubicin behaves as an intercalator during DNA replication. *(A)* Plasmid DNA was replicated in the presence of [α-^32^P]dATP and vehicle control, aclarubicin, or doxorubicin. *(B)* Samples from (A) were separated on a native agarose gel and visualized by autoradiography. *(C)* Quantification of supercoiled circular monomers from (B). Mean ± SD, n=3 independent experiments. See also Supplemental Figure 5A. *(D)* Plasmid DNA was replicated in the presence of vehicle control or doxorubicin. In parallel, plasmid DNA was incubated with LacR, which bound to a *lacO* array within the plasmid. After 20 minutes, when forks were localized either side of the LacR array, vehicle control or doxorubicin were added, along with IPTG to restart DNA replication. Plasmids were purified and digested with XmnI, which cuts the plasmid once and allows replication fork structures (‘DYs’) and fully replicated molecules (‘Lins’) to be identified. *(E)* Samples from (D) were separated on a native agarose gel and visualized by autoradiography. *(F)* Quantification of fork merger from (E) lanes 1-8. Mean ± SD, n=3 independent experiments. *(G)* Quantification of fork merger from (E) lanes 9-17. Mean ± SD, n=3 independent experiments.

Intercalators bind double-stranded DNA, but DNA synthesis takes place at single-stranded DNA unwound by the replicative helicase (Bell and Labib 2016). Thus, we reasoned that if doxorubicin inhibits replication by intercalating into DNA it would have to do this by intercalating into the unreplicated DNA and inhibiting DNA unwinding. In this case, doxorubicin should preferentially inhibit earlier stages of replication (‘initiation’) due to there being more DNA template. To test this, we administered doxorubicin at the onset of replication or just prior to completion of DNA synthesis (‘termination’). To this end, we added doxorubicin either at the onset of DNA replication or after localizing replication forks to a LacR-bound array of *lacO* sequences (Fig. 5D), which functions as an efficient replication barrier (Dewar et al. 2015). In both cases we measured fork merger, which represents the point at which the final stretch of DNA is unwound, as a read-out for replication fork progression (Fig. 5D). Addition of doxorubicin at the onset of replication completely blocked DNA replication (Fig. 5E, lanes 5-8) evidenced by the complete lack of radioactive signal compared to the control (Fig. 5E, lanes 1-4). Accordingly, fork merger was readily detected in the control but not following doxorubicin addition (Fig. 5F). Surprisingly, the same concentration of doxorubicin that completely blocked replication initiation had no discernable effect when added prior to termination (Fig. 5E, lanes 10-17, 5G). Thus, doxorubicin preferentially inhibits initiation of DNA replication.

To test whether doxorubicin could also inhibit replication fork progression (‘elongation’), in addition to initiation, of DNA synthesis we measured fork merger in the presence of increasing concentrations of doxorubicin. When the concentration of doxorubicin was increased 2-fold, to 1.4 µM, fork merger was strongly inhibited (Supplemental Fig. S5C, lanes 10-13, S5D), while 4-fold higher completely blocked fork merger (Supplemental Fig. S5C, lanes 14-17, S5D). This demonstrated that increasing concentrations of doxorubicin inhibit either elongation or termination of DNA synthesis. To distinguish between these possibilities, we tested the ability of doxorubicin to inhibit fork progression through a ∼750 base pair *lacO* array compared to a ∼1500 base pair *lacO* array (Supplemental Fig. S5E). If doxorubicin primarily inhibits termination, then the extent of inhibition by doxorubicin should be the same regardless of the amount of DNA synthesized in the presence of the drug. Alternatively, if doxorubicin primarily inhibits elongation, then inhibition should be approximately doubled according if twice as much DNA synthesis occurs in the presence of the drug. Doxorubicin inhibited fork merger within the ∼750 base pair array by ∼5 minutes (Supplemental Fig. S5F, H) and for the ∼1500 bp array this increased to ∼12 minutes (Supplemental Fig. S5G,I). Thus, these data show that higher concentrations of doxorubicin preferentially inhibit elongation, rather than termination, consistent with there being very little unreplicated DNA during termination. Overall, we conclude that DNA intercalation by doxorubicin inhibits both initiation and elongation of DNA synthesis but preferentially inhibits initiation.

### Etoposide and doxorubicin inhibit replication fork progression through different mechanisms in mammalian cells

Doxorubicin toxicity is largely dependent on Top2α, which is thought to primarily act during DNA replication (Burgess et al. 2008; Olivieri et al. 2020). We were therefore surprised by the strong Top2α-independent inhibition of replication by doxorubicin. To test whether our drug stocks reproduced the previously reported cell killing effects we examined the viability of U2OS cells exposed to etoposide and doxorubicin in the presence or absence of Top2α. Cells were treated with control or Top2α siRNA, then etoposide or doxorubicin for 24 hours, before the drug was washed out to allow for colony formation (Fig. 6A). The concentrations of etoposide and doxorubicin we used resulted in near-complete loss of viability for cells treated with control siRNA (Fig. 6B). Importantly, siRNA knock down of Top2α restored most of the loss of viability in both cases (Fig. 6B). Thus, cell killing by the exact same drug stocks we used in our experiments (Fig. 1-5) is largely Top2α-dependent, as previously reported.

**Figure 6.**
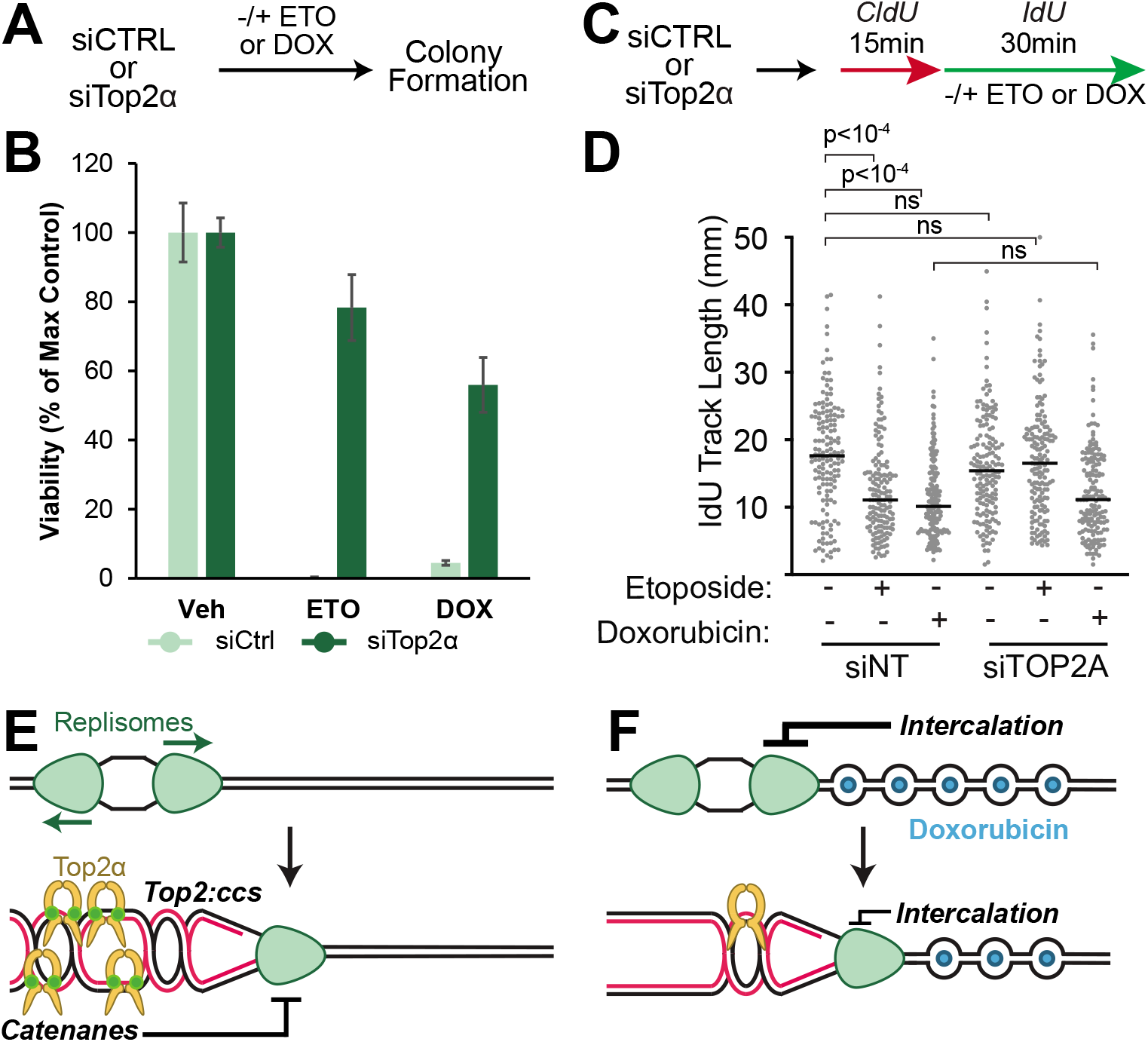
Etoposide and doxorubicin inhibit mammalian DNA replication through distinct mechanisms. *(A)* U2OS cells were treated with siRNA targeting Top2α (‘siTop2α’) or control siRNA (‘siCtrl’), then treated with etoposide or doxorubicin for 24 hours. Drugs were washed out and cells were allowed to form colonies for 10 days before being counted. *(B)* Quantification of cell viability from (A). Mean ± SD, n=3 independent experiments. *(C)* HCT116 cells were treated with siRNA targeting Top2α (‘siTop2α’) or control siRNA (‘siCtrl’). Cells were then labeled with CldU and IdU and treated with etoposide or doxorubicin for the indicated times prior to DNA combing. *(D)* Median IdU track length of individual CldU-positive fibers from (C). p values were derived from an ANOVA with Dunn’s multiple comparisons post-test in all panels. *(E)* Etoposide traps Top2α on newly replicated DNA behind replication forks, preventing catenane resolution. This leads to an accumulation of topological stress that stalls the replication fork *(F)* Doxorubicin intercalates parental DNA ahead of the fork, slowing replication fork progression. Earlier stages of replication are inhibited more than later stages of replication due to the amount of DNA available in front of the fork.

We next wanted to test whether the effects of etoposide and doxorubicin that we observed in *Xenopus* egg extracts also took place in cells. We took advantage of the fact that replication fork progression is inhibited by both etoposide (Fig. 4) and doxorubicin (Supplemental Fig. S5B-D) and this depends on Top2α for etoposide but not for doxorubicin (Fig. 3A-D). To test whether this is also true in mammalian cells, we treated HCT116 cells with either control or Top2α siRNA, then performed DNA combing to monitor the effects of etoposide and doxorubicin on replication fork progression (Fig. 6C). We found that both etoposide and doxorubicin inhibited fork progression in cells treated with control siRNA (Fig. 6D) as previously reported (Zellweger et al. 2015). When Top2α was silenced, there was no significant impact on fork progression (Fig. 6D), consistent with our previous observation that loss of Top2α has little impact on total DNA synthesis (Heintzman et al. 2019). Importantly, treatment with Top2α siRNA rescued the replication slowing induced by etoposide but doxorubicin still inhibited DNA replication (Fig. 6D), consistent with our results in extracts (Fig. 3A-D). We conclude that etoposide inhibits fork progression in a Top2α-dependent manner while doxorubicin inhibits DNA replication in a Top2α-independent manner, both in *Xenopus* egg extracts and mammalian cells.

## Discussion

We have identified new mechanisms of replication inhibition by topoisomerase II (Top2) poisons, which can be explained by the following models. Etoposide traps Top2α behind replication forks, which leads to fork stalling, due to accumulation of topological stress (Figure 6E). Doxorubicin acts as a DNA intercalator and inhibits all stages of replication by inhibiting unwinding of the replicative helicase (Figure 6F). These points are discussed further below.

### Replication inhibition by etoposide

In our model (Figure 6E), etoposide traps Top2α behind forks, which prevents catenanes from being resolved and stalls replication forks. This model is consistent with Top2 poisons acting early in the Top2 catalytic cycle (Pommier et al. 2016; Vann et al. 2021). Interestingly, etoposide interferes with fork progression at all stages of DNA synthesis, while our previous work showed that loss of Top2α stalls forks specifically during termination (Heintzman et al. 2019). Depletion of Top2α essentially blocks catenane resolution does not inhibit DNA replication until termination (Heintzman et al. 2019), so it is not clear why preventing catenane resolution by etoposide would inhibit replication at earlier stages. One possibility is that etoposide traps catenanes close behind replication forks, which exacerbates the deleterious effects of catenanes. Accordingly, partial depletion of Top2α reduced the impact of etoposide and resulted in fork stalling only during termination (Supplemental Fig. 4A-D). Additionally, replacement of Top2α with a catalytic dead allele, which would also be expected to trap catenanes behind forks, also led to fork stalling prior to termination (Heintzman et al. 2019). It will be important to definitively determine whether catenane trapping by etoposide causes replication fork stalling. If so, proteins that limit diffusion of topological stress, such as cohesin (Minchell et al. 2020), might regulate the response to etoposide. Additionally, targeting cell cycle regulatory mechanisms (Morafraile et al. 2019) and replication proteins (Schalbetter et al. 2015) that normally limit catenane generation may enhance the cytotoxicity of etoposide.

Top2β is present in cells and can readily relax DNA supercoils (Austin et al. 2018). If supercoils are formed during vertebrate DNA replication Top2β should still be trapped on them by etoposide, even following Top2α inactivation, and this would be expected to slow replication. It is therefore surprising that in cells inactivation of Top2α is sufficient to eliminate replication slowing by etoposide. This suggests that DNA supercoil formation is rare during vertebrate DNA replication. This would be consistent with previous observations in *Xenopus* egg extracts that Top2α, which can act on both supercoils and catenanes, primarily acts behind replication forks (Lucas et al. 2001; Heintzman et al. 2019) suggesting that supercoils are rarely formed. In the future it will be important to determine how the frequency of supercoils compared to catenanes is determined during vertebrate DNA replication.

### Replication inhibition by doxorubicin

Our model proposes that doxorubicin inhibits DNA unwinding by intercalating into the parental DNA (Figure 6F). The simplest explanation is that intercalation stabilizes duplex DNA and forms a barrier to unwinding by the replicative helicase. In support of this model, doxorubicin preferentially inhibits earlier stages of DNA synthesis, consistent with its activity being dependent on unreplicated DNA. Although doxorubicin has been reported to induce oxidative DNA damage and DNA cross-links (Doroshow et al. 2001; Coldwell et al. 2008) we do not observe any alterations in DNA structures suggesting that DNA damage does not take place. Additionally, replication slowing is unlikely to be an indirect consequence of histone eviction by doxorubicin (Pang et al. 2013; Yang et al. 2013) because histones constrain, rather than promote, fork progression (Kurat et al. 2017). The intercalation activity of doxorubicin that inhibits DNA replication would be expected to broadly inhibit DNA metabolic processes that generate the substrates for Top2. Thus, it will be important to test whether modified forms of doxorubicin that are weaker intercalators can exhibit more potent cell killing.

### Implications for cell killing

It is surprising that etoposide and doxorubicin behave differently during DNA replication given that the both exhibit Top2α-dependent cytotoxicity. Since Top2α functions during multiple cellular processes (Pommier et al. 2016; Vann et al. 2021) our favored model is that doxorubicin and etoposide target different aspects of chromosome metabolism, such as during transcription or chromosome organization. In line with this idea, it has been recently shown that G-quadraplex stabilizing agents CX-5461 and pyridostatin both act as Top2 poisons at sites of transcription by stabilizing G-quadruplex structures (Olivieri et al. 2020; Bossaert et al. 2021). Doxorubicin was also reported to stabilize G-quadruplex structures (Scaglioni et al. 2016), suggesting that it might work in the same way. However, there are other potential explanations for these results. For example, doxorubicin is much less efficient than etoposide at inducing DNA breaks (Binaschi et al. 1990) so it is possible that doxorubicin acts as a pro-drug and needs to be modified to become an active Top2 poison. This would induce cell killing in long-term cell viability assays but would be missed in the short-term time course replication assays we performed. In the future it will be important to determine exactly how etoposide and doxorubicin elicit Top2-dependent cell killing.

## Methods

### Xenopus egg extracts

*Xenopus* egg extracts were prepared from *Xenopus laevis* wild-type male and female frogs (Nasco) as previously described (Lebofsky et al. 2009). Animal care protocols were approved by the Vanderbilt Division of Animal Care (DAC) and Institutional Animal Care and Use Committee (IACUC).

### Plasmid Construction and Preparation

The construction of pJD85(p[*lacO*x8]), pJD88(p[lacOx16]), pJD100(p[*lacO*x48]) and pJD152(p[*lacO*x16]) was described previously (Dewar et al. 2015). To create pJD90(p[*lacO*x24]), the BsrGI/BsiWI DNA fragment from pJD85(p[*lacO*x8]) was cloned into pJD88(p[lacOx16]) after it had been digested by BsrGI/BsiWI. pJD90(p[*lacO*x24]) and pJD100(p[*lacO*x48]) were used in Supplemental Fig. 5E-I. All other experiments used JD152(p[*lacO*x16]) as a template for replication.

### DNA Replication in Xenopus egg extracts

DNA replication was performed as previously described (Dewar et al. 2015). High Speed Supernatant (‘HSS’) was incubated with nocodazole (3 ng/μl) and ATP regenerating system (‘ARS,’ 20 mM phosphocreatine, 2 mM ATP and 5 ng/μl phosphokinase) for five minutes at room temperature. When indicated, LacR protein was bound to *lacO* repeats on plasmid DNA for one hour at room temperature before licensing DNA. To license plasmid DNA, one volume of ‘licensing mix’ was prepared by incubating plasmid DNA (final concentration of DNA in licensing mix was 12.3 ng/μl for most experiments; final concentration of DNA in licensing mix was 7.32 ng/μl in Supplemental Fig. 5E-I) with HSS at room temperature for 30 minutes. NucleoPlasmic Extract (‘NPE’) was supplemented with ARS, DTT (final concentration: 2 mM), NPE. To initiate replication, 2 volumes of NPE mix were added to 1 volume of licensing mix. If applicable, replication forks that were stalled at a LacR-bound *lacO* array were released by IPTG addition as described previously (Dewar et al. 2015). Samples were withdrawn into Replication Stop Solution (8 mM EDTA, 0.13% phosphoric acid, 10% ficoll, 5% SDS, 0.2% bromophenol blue, 80 mM Tris, pH 8) then vigorously mixed, and treated with Proteinase K (929 ng/μl) before analysis via native agarose gel electrophoresis. Etoposide and Doxorubicin were dissolved in DMSO and used at the concentrations. p97 inhibitor (NMS-873, p97-i) was dissolved in DMSO and used at a final concentration of 200 µM. Etoposide, doxorubicin, and p97-i were used with a final concentration of 4% (v/v) DMSO in the reaction. Caffeine was dissolved in water and used at a final concentration of 5 mM. ATR inhibitor (VE-821, ATRi) was dissolved in water and used at a final concentration of 10 μM.

### Breaks Assay

For breaks assays, samples were withdrawn into Extraction Stop Solution (0.5% SDS, 25 mM EDTA, 50 mM Tris-HCl, pH 7.5), then treated with RNase A (334 ng/μl) for 30 minutes at 37°C, followed by Proteinase K (769 ng/μl) for 1 hour at 37°C. Samples were then purified by phenol-chloroform extraction and digested with XmnI restriction enzyme (New England BioLabs) as previously described (Dewar et al. 2015). To identify DNA fragment sizes, 100 bp and 1 kb DNA ladders from New England BioLabs were radiolabeled with T4 PNK then separated alongside XmnI-digested samples. Breaks were quantified by measuring radioactive signal between 500 and ∼1500 bp for the ∼3100 bp plasmid template.

### DNA-Protein Crosslink Detection Assay

DNA-Protein Crosslinks (DPCs) were detected as previously described (Duxin et al. 2014). Briefly, replication assay samples were sampled in twice the normal volume of Replication Stop, then split into two. One half of the sample was treated with Proteinase K (889 ng/μl) as described above and the other half was treated with filtered water. DNA molecules were visualized via native agarose gel electrophoresis. DPCs were quantified by calculating the percentage of radioactive signal above the θ to the wells of the gel then subtracting the signal in the +ProtK condition from the -ProtK condition.

### Protein Purification

Biotinylated LacR protein was expressed in *Escherichia coli* bacteria cells and purified as described previously (Dewar et al. 2015).

### Antibodies

Antibodies targeting *Xenopus* CDC45, MCM6 and TOP2α were the same as those previously described (Heintzman et al. 2019). The following antibodies were used to detect human proteins: TOP2α (Bethyl, Cat# A300-054A), TOP2β (Topogen, Cat# tg2010-3).

### Immunodepletions

Top2α was immunodepleted from extract as previously described (Heintzman et al. 2019). Depleted extracts were collected and used for DNA replication (as above). For rescue experiments (Supplemental Fig. 3), human recombinant Top2α was added at a final concentration of 20.78 μg/ml. For the experiments in Fig 3E-H, plasmid DNA was first allowed to replicate in Top2α depleted extract to allow for catenanes to form. Undepleted extract was then diluted with two volumes 1X ELB to make 33% undepleted NPE. Then at 60 minutes, the extract was split into equal volumes and treated with drug as indicated. After 30 seconds of drug incubation with the immunodepleted extract, 1/16^th^ volume of 33% undepleted NPE was added to add back *Xenopus* Top2α protein to ∼6% of undepleted levels.

### Analysis of chromatin-bound proteins and Western blotting

Plasmid pull downs and Western blotting in *Xenopus* egg extracts were performed as previously described (Dewar et al. 2017). Western blotting of mammalian cell lysates were also performed as previously described (Mehta et al. 2020).

### siRNA

The following siRNAs were used: AllStars Negative Control siRNA (Qiagen, Cat#1027281) and TOP2A (Dharmacon, Cat# J-004239-06-0002, J-004239-07-0002, J-004239-08-0002, and J-004239-09-0002)

### Cell Viability Assay

U2OS cells were transfected with 40 pmol siRNA for 24 hours and plated 24 hours later (300 cells/ 60 mm dish). Cells were then treated with drug (Etoposide final concentration: 1 μM; doxorubicin final concentration: 16 nM) 24 hours later for a total incubation of 24 hours. Drugs were washed out and cells were then allowed to recover in fresh media for 10 days. Colonies were then stained with methylene blue (48% methanol, 2% methylene blue, 50% water) and counted.

### DNA Molecular Combing

Cells were labeled with 20mM CldU (Sigma, C6891) followed by 100mM IdU (Sigma, l7125), for the times indicated, with or without 20 µM Etoposide or 10 µM Doxorubicin added with the second nucleoside analog treatment. Approximately 600,000 cells were embedded in 1.5% low-melting agarose plugs in PBS and were digested overnight in 0.1% Sarkosyl, 2mg/ml proteinase K, and 50 mM EDTA, pH 8.0 at 50° C. Plugs were washed in TE pH 8.0, transferred to 100mM MES pH5.7, melted at 68° C and digested with 1.5 units b-Agarase overnight at 42°C. DNA was combed onto silanized coverslips (Microsurfaces, Inc., Custom) using a Genomic Vision combing apparatus. The DNA was stained with antibodies that recognize IdU and CldU (Abcam Cat#ab6326, BD Cat#347580) for 1 hour, washed in PBS, and probed with secondary antibodies for 30 minutes. Images were obtained using a 40X oil objective (Nikon Eclipse Ti). Analysis of fiber lengths performed using Nikon Elements software. Statistical analyses were performed with Graphpad Prism 9.

## Acknowledgements

JMD was supported by NIH grant R35GM128696. NO was supported by R01GM126363. DC was supported by R01CA239161. KPM was supported by F32GM136096. We thank Stedman Stephens for experimental assistance and Stephanie Medina for sharing unpublished observations.

---

**Supplemental Figure 1.**
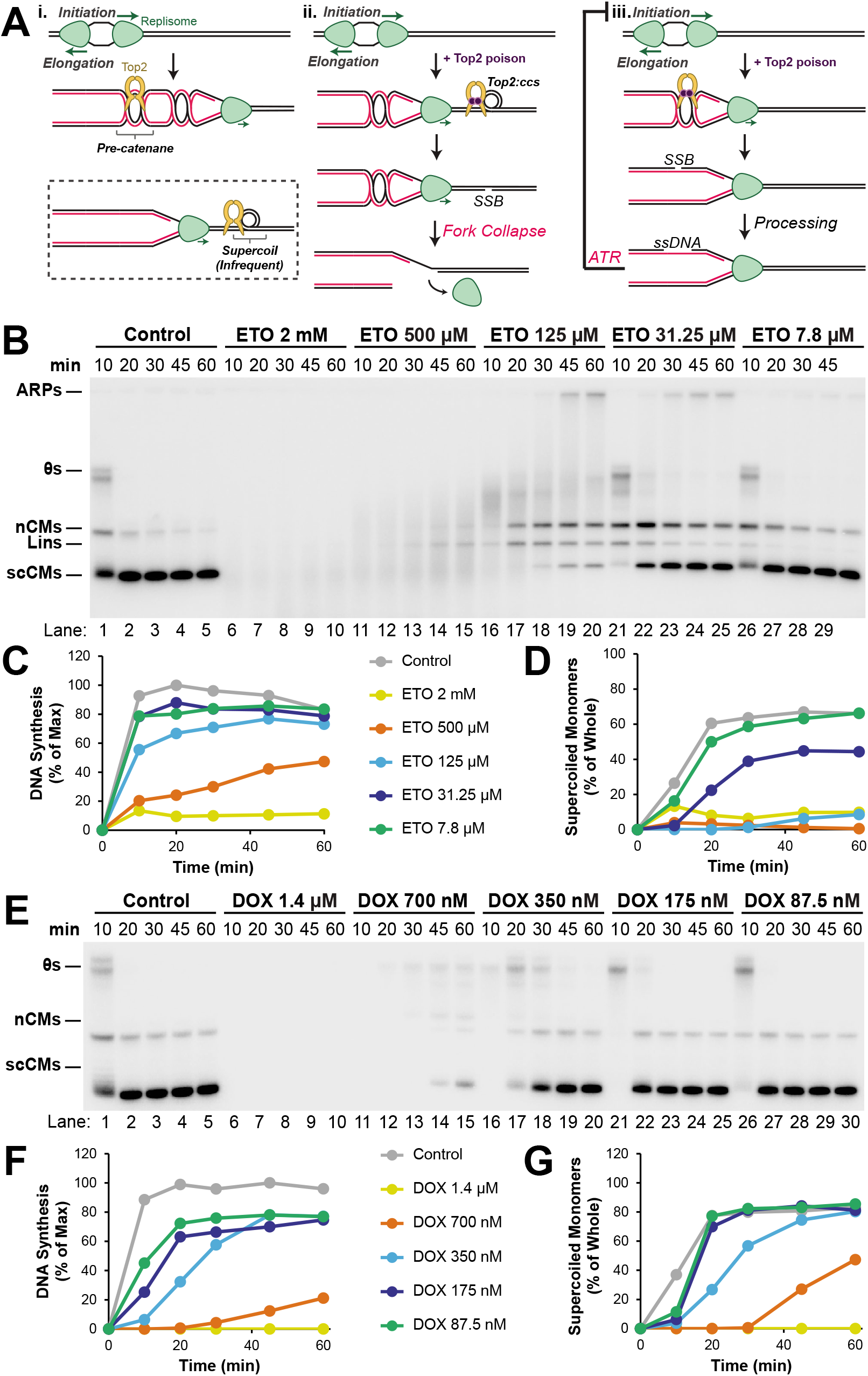
Effects of topoisomerase poisons on DNA synthesis. (A) Models for how Top2 poisons impact DNA replication. (i) During normal DNA replication, two replisomes activate and begin unwinding the DNA, moving in opposite directions (‘Initiation’). As DNA is replicated (‘Elongation’), pre-catenanes form behind the replication fork on newly-synthesized DNA, which is recognized and resolved by Top2. Supercoils are infrequent during vertebrate DNA replication (Lucas et al. 2001; Heintzman et al. 2019) so catenanes are likely to be the main substrate for Top2 during DNA synthesis. (ii) Top2 poisons have been proposed to trap Top2 ahead of the fork on supercoils, leading to DNA breaks. If the replication fork encounters a DNA break, the fork will collapse and replisome proteins will passively dissociate from DNA (Vrtis et al. 2021). (iii) Top2 poisons can induce single-strand DNA breaks which are expanded to gaps and activate the ATR checkpoint to inhibit replication initiation (Costanzo et al. 2003). *(B)* Plasmid DNA was replicated as in Figure 1A but with a wide range of concentrations of etoposide. *(C)* Quantification of total DNA synthesis from (B). *(D)* Quantification of supercoiled circular monomers (‘scCMs’) from (B). *(E)* Plasmid DNA was replicated as in Figure 1A but with a wide range of concentrations of doxorubicin. *(F)* Quantification of total DNA synthesis from (E). *(G)* Quantification of scCMs from (E).

**Supplemental Figure 2.**
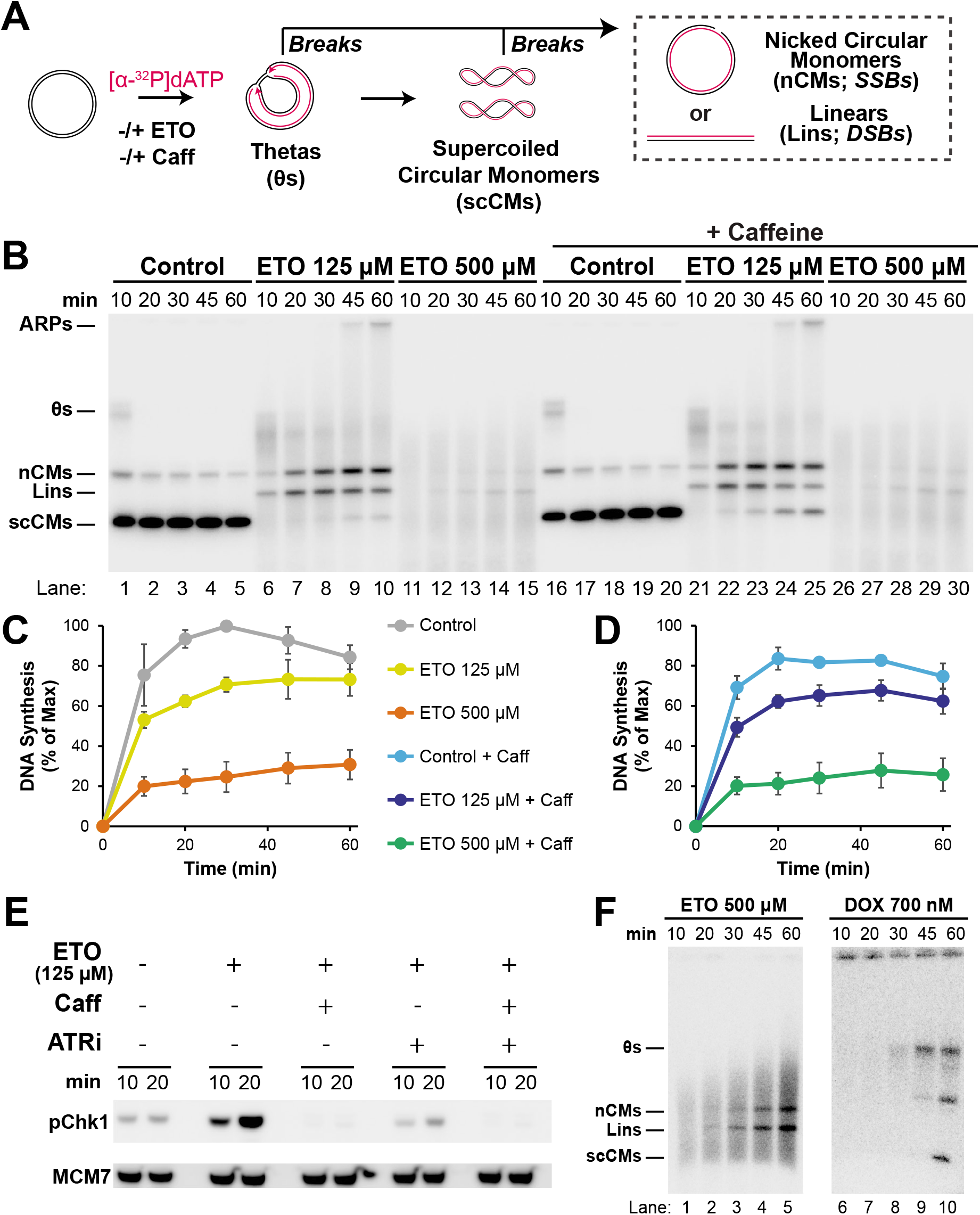
Etoposide inhibits replication independent of ATR activation. *(A)* To test whether the effects of etoposide on DNA replication are due to ATR signaling, as previously described (Costanzo et al. 2003), plasmid DNA was replicated in the presence or absence of etoposide (‘ETO’) and caffeine (‘Caff’). *(B)* Samples from (A) were separated on a native agarose gel and visualized by autoradiography. *(C)* Quantification of total DNA synthesis from (B) lanes 1-15. Mean ± SD, n=3 independent experiments. Etoposide led to a dose-dependent inhibition of DNA synthesis, as in Fig 1. *(D)* Quantification of total DNA synthesis from (B) lanes 16-30. Mean ± SD, n=3 independent experiments. Etoposide led to a dose-dependent inhibition of DNA synthesis, as in (C), even though caffeine should block ATR activation. *(E)* To confirm that the ATR checkpoint was activated in (B) and that this was blocked by caffeine, DNA was replicated as in (A) in the presence or absence of ATR inhibitor (VE-821, ‘ATR-i’). Western blotting was performed to measure the abundance of phosphorylated Chk1 (‘pChk1’) as a read-out for ATR activation. MCM7 was detected as a loading control. Etoposide induced a strong pChk1 signal that was diminished by ATRi, indicating that ATR was activated in response to etoposide, as previously described (Costanzo et al. 2003). Importantly, caffeine completed blocked pChk1 signal indicating that caffeine blocked ATR activation. Thus, ATR signaling was not responsible for the replication slowing in (B)-(C). *(F)* Overexposure of indicated conditions from Fig. 1B.

**Supplemental Figure 3.**
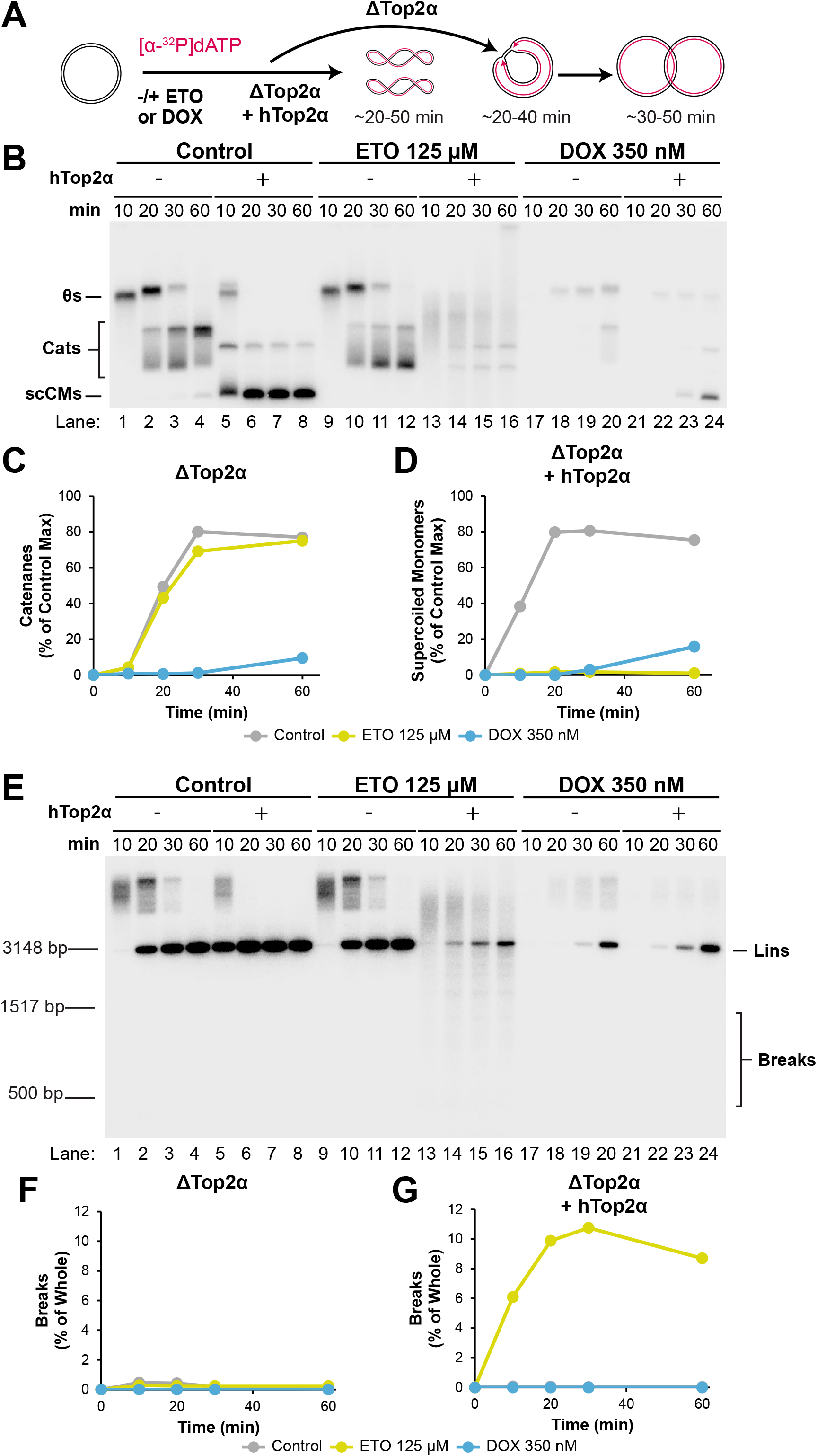
Doxorubicin does not act as a topoisomerase poison against human Top2α during DNA replication. *(A)* To test whether doxorubicin acted as a topoisomerase poison against human Top2α (hTop2α) the endogenous Top2α was depleted from extracts, which were then used to replicate DNA in the presence of either buffer control or hTop2α. Vehicle control, etoposide, or doxorubicin were also added, as indicated. In the buffer control (ΔTop2α) the final replication products should be catenanes (‘Cats’) due to the lack of Top2 activity. In the hTop2α condition (ΔTop2α+hTop2α) and the final replication products should be supercoiled circular monomers (‘scCMs’) as hTop2α rescues the depletion (Heintzman et al. 2019). *(B)* Samples from (A) were separated on a native agarose gel and visualized by autoradiography. *(C)* Quantification of catenanes in ΔTop2α conditions from (B) lanes 1-4, 9-12, and 17-20. Doxorubicin, but not etoposide, slows catenane formation as in Fig. 3D. *(D)* Quantification of supercoiled monomers in ΔTop2α + hTop2α conditions from (B) lanes 6-8, 13-16, and 21-24. Both etoposide and doxorubicin slow replication, indicating that etoposide, but not doxorubicin, inhibits replication in a hTop2α-dependent manner. *(E)* To test the ability of etoposide and doxorubicin to induce breaks by hTop2α, samples from (A) were digested by XmnI, separated on a native agarose gel, and visualized by autoradiography. *(F)* Quantification of breaks in ΔTop2α conditions from (E) lanes 1-4, 9-12, and 17-20. In the absence of Top2α breaks are not readily detected. *(G)* Quantification of breaks in ΔTop2α + hTop2α conditions from (E) lanes 6-8, 13-16, and 21-24. In the presence of hTop2α etoposide induces breaks, but doxorubicin does not, as observed in Fig. 2C. Overall, etoposide behaves as a Top2 poison in the presence of hTop2α but doxorubicin does not, as observed for *Xenopus* Top2α in undepleted extracts (Figs. 2-3).

**Supplemental Figure 4.**
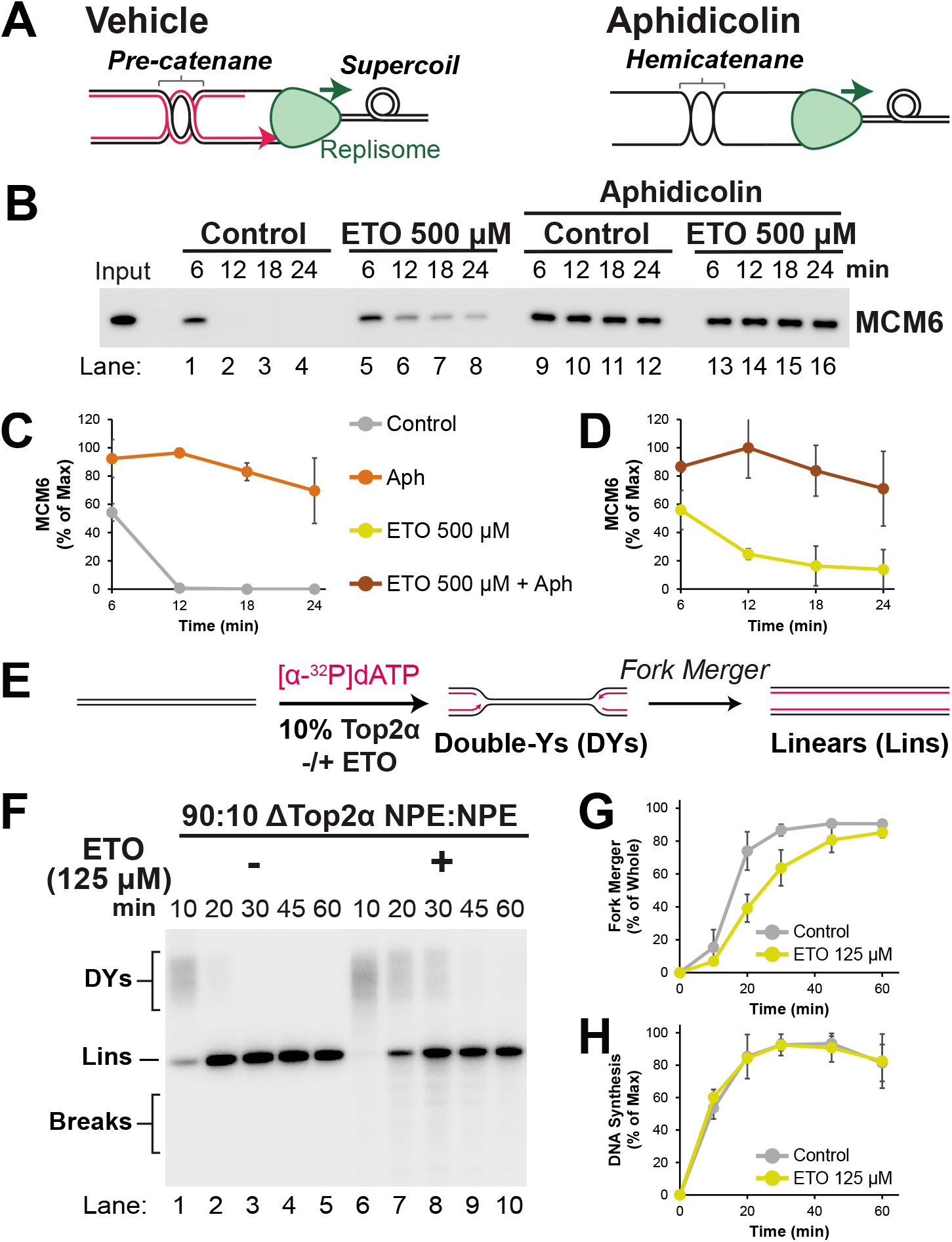
Etoposide inhibits DNA replication by trapping Top2 on newly-replicated DNA. *(A)* Cartoon depicting the DNA structures formed by DNA replication in the presence of aphidicolin or a vehicle control. In the vehicle control both supercoils and catenanes can form. However, upon aphidicolin treatment, DNA synthesis is blocked so only hemicatenanes, and not catenanes, can be formed behind the fork. Note that in the presence of aphidicolin extensive DNA unwinding and supercoil generation takes place (Walter and Newport 2000). *(B)* Samples from Fig. 4G were analyzed by Western blotting to detect MCM6. *(C)* Quantification of (B) lanes 1-4 and 9-12. Mean ± SD, n=3 independent experiments. *(C)* Quantification of MCM6 from (B) lanes 5-8 and 13-16. Mean ± SD, n=3 independent experiments. *(E)* To test whether etoposide could also cause termination defects, as observed following loss of Top2 (Heintzman et al. 2019), replication was performed with reduced amounts of Top2 in the absence or presence of etoposide. Plasmid DNA was replicated in extracts where Top2α was reduced 10-fold by mixing mock- and Top2α-immunodepleted extracts, which was sufficient to prevent fork stalling (Heintzman et al. 2019). Purified DNA was then digested with XmnI so that replication forks (double-Ys) and replicated molecules (linears) could be identified and fork merger could be measured. *(F)* Samples from (E) were separated on a native agarose gel and visualized by autoradiography. *(G)* Quantification of fork merger from (F). Note that breaks (broken molecules) were excluded from the analysis as these would cause the rate of fork merger to be under-estimated. Mean ± SD, n=3 independent experiments. *(H)* Quantification of total DNA synthesis from (F). Mean ± SD, n=3 independent experiments.

**Supplemental Figure 5.**
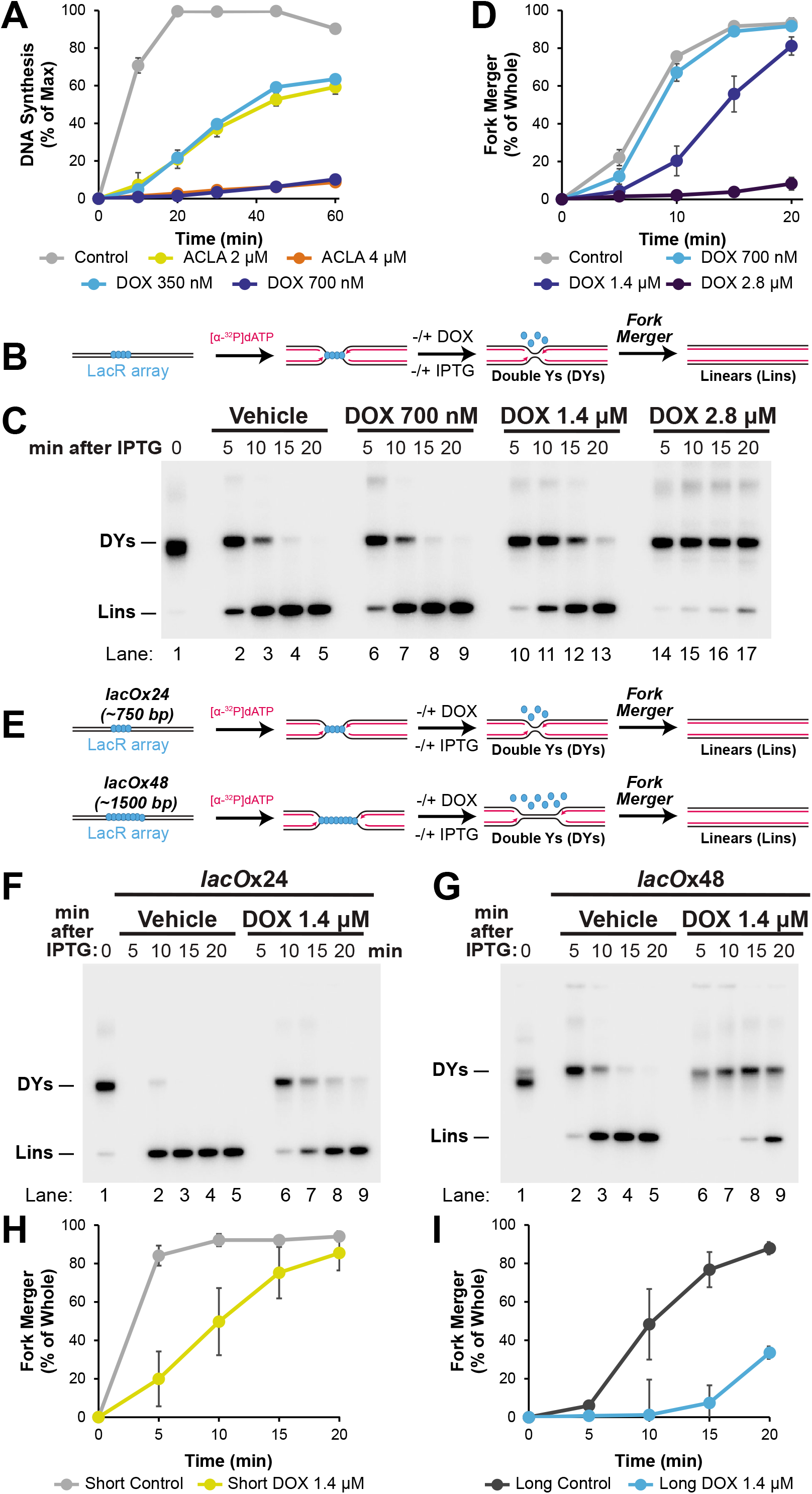
Doxorubicin inhibits replication through intercalation of unreplicated DNA. *(A)* Quantification of total DNA synthesis from Fig. 5B. Mean ± SD, n=3 independent experiments. *(B)* Plasmid DNA harboring a LacR array was replicated for 19 minutes to localize replication forks each side of the barrier. Increasing concentrations of doxorubicin were added one minute before IPTG was added to release the barrier and restart replication. Purified replication intermediates were digested with XmnI so that replication fork structures (double-Ys) and fully replicated molecules (linears) could be identified. *(C)* Samples from (B) were separated on a native agarose gel and visualized by autoradiography. *(D)* Quantification of fork merger from (C). Mean ± SD, n=3 independent experiments. *(E)* Replication was performed as in (B) but with a *lacO*x24 (∼750 bp) *and lacO*x48 (∼1500 bp) LacR array and a single concentration of doxorubicin (1.4 µM). *(F) lacO*x24 array samples from (E) were separated on a native agarose gel and visualized by autoradiography. *(G) lacO*x48 array samples from (E) were separated on a native agarose gel and visualized by autoradiography. *(H)* Quantification of fork merger from (F). Mean ± SD, n=3 independent experiments. *(I)* Quantification of fork merger from (G). Mean ± SD, n=3 independent experiments.

**Supplemental Figure 6.**
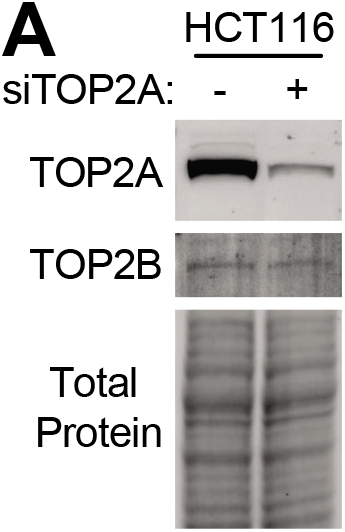
Top2α and Top2β levels following siRNA of TOP2A. *(A)* Cells from Figure. 6C were lysed and analyzed by Western blotting. siTOP2A dramatically reduces levels of Top2α protein but not Top2β.

## Notes

### Competing Interest Statement

The authors have declared no competing interest.

### Summary of Updates

The manuscript has been revised to include funding information in the acknowledgements.

